# PSGL-1 inhibits the virion incorporation of SARS-CoV and SARS-CoV-2 spike glycoproteins and impairs virus attachment and infectivity

**DOI:** 10.1101/2020.05.01.073387

**Authors:** Sijia He, Abdul A. Waheed, Brian Hetrick, Deemah Dabbagh, Ivan V. Akhrymuk, Kylene Kehn-Hall, Eric O. Freed, Yuntao Wu

**Affiliations:** National Center for Biodefense and Infectious Diseases, School of Systems Biology, George Mason University, Manassas, VA 20110, USA; Virus-Cell Interaction Section, HIV Dynamics and Replication Program, Center for Cancer Research, National Cancer Institute-Frederick, Frederick, MD 21702, USA

**Author notes:** These authors contributed equally to this work (Sijia He, Abdul A. Waheed).

**Keywords:** Coronavirus, cellular antiviral factor, pseudovirion

## Abstract

P-selectin glycoprotein ligand-1 (PSGL-1) is a cell surface glycoprotein that binds to P-, E-, and L-selectins to mediate the tethering and rolling of immune cells on the surface of the endothelium for cell migration into inflamed tissues. PSGL-1 has been identified as an interferon-γ (INF-γ)-regulated factor that restricts HIV-1 infectivity, and has recently been found to possess broad-spectrum antiviral activities. Here we report that the expression of PSGL-1 in virus-producing cells impairs the incorporation of SARS-CoV and SARS-CoV-2 spike (S) glycoproteins into pseudovirions and blocks virus attachment and infection of target cells. These findings suggest that PSGL-1 may potentially inhibit coronavirus replication in PSGL-1^+^ cells.

---

The ongoing coronavirus disease 2019 (COVID-19) is a global pandemic afflicting more than 10 million people in over 200 countries and territories, resulting in more than 500,000 deaths as of June 30th, 2020. Currently, there are no effective treatments or vaccines. Understanding virus-host interactions is critical for developing novel therapeutics and vaccines. P-selectin glycoprotein ligand-1 (PSGL-1, also known as SELPLG or CD162) is a human protein recently identified to possess broad-spectrum antiviral activity (1). PSGL-1 binds to the selectin family of proteins, P-, E-, and L-selectin (2), and mediates immune cell tethering and rolling on the surface of endothelium to promote cell migration into inflamed tissues (3). In the context of viral infection, PSGL-1 has been identified as an IFN-γ-regulated inhibitory factor involved in blocking HIV-1 infectivity (4), and was recently found to possess broad-spectrum antiviral activity (1), blocking viral infections through steric hindrance of particle attachment to target cells (1, 5).

The coronavirus spike (S) glycoproteins play an essential role in viral entry by binding the cell-surface receptor on target cells and mediating the fusion between viral and cellular membranes during virus entry (6). The S protein is also the target of neutralizing antibodies generated by the infected host. Because of its central role in virus infection and adaptive immunity, the S protein is a prime target for the development of antiviral therapeutics and vaccines. In addition to the adaptive arm of the host immune response, viral infections trigger an innate immune response, largely induced by IFN, that sets up an antiviral state. Hundreds of IFN-stimulated genes (ISGs) are induced by viral infection (7). While the role of some ISGs in blocking the replication of particular viruses has been well established, the vast majority of ISGs have not been characterized. Because of the significance of host innate immunity in viral transmission and replication within and between hosts, there is an unmet need to understand these antiviral inhibitory factors in detail.

Previous studies have demonstrated that PSGL-1 can be incorporated into HIV-1 virions, and its virion incorporation subsequently blocks viral infectivity (1, 5). To investigate the ability of PSGL-1 to restrict coronavirus infection, we first established a lentiviral vector-based coronavirus pseudovirus infection system (8), in which the S proteins from either SARS-CoV or SARS-CoV-2 were used to pseudotype lentiviral particles (**Fig. 1A**). Using this system, we assembled particles in the presence or absence of PSGL-1 (1), and then used the particles to infect target Vero and Calu-3 cells, which endogenously express the primary SARS-CoV and CoV-2 receptor, angiotensin converting enzyme 2 (ACE2) (9, 10). The expression of PSGL-1 in viral producer cells had a minor (~ two-fold) effect on the release of SARS-CoV and –CoV-2 pseudovirions (**Fig. 1B and 1C**), consistent with the previous finding that PSGL-1 expression has minimal effects on viral release (1). However, the infectivity of PSGL-1-imprinted SARS-CoV particles was completely abrogated in Vero cells (**Fig. 1D**), demonstrating the ability of PSGL-1 to block the infectivity of SARS-CoV S-bearing virions.

**Fig. 1.**
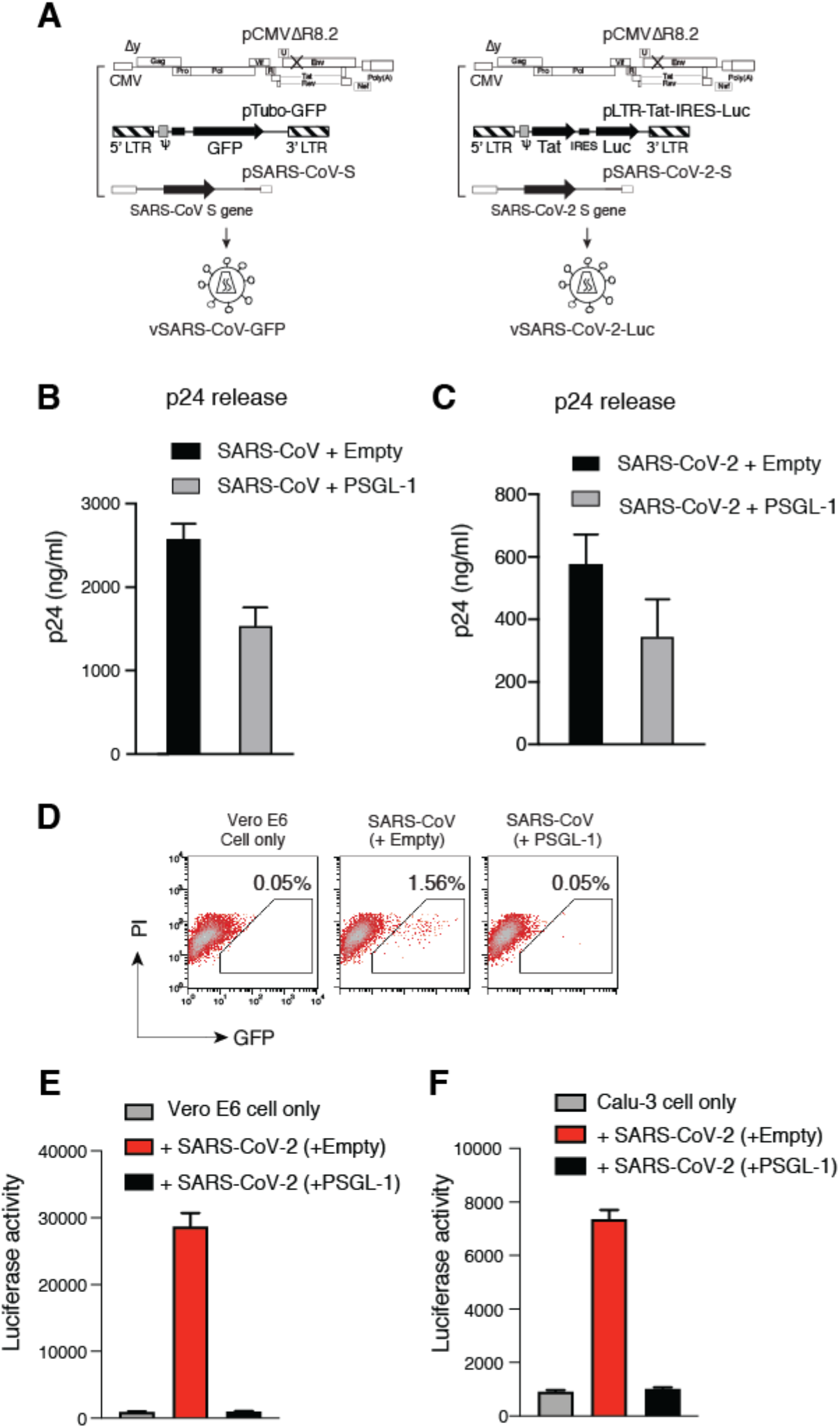
PSGL-1 inactivates the infectivity of SARS-CoV and SARS-CoV2 pseudoviruses. **A)** Schematic of the assembly of lentiviral particles pseudotyped with the S proteins of SARS-CoV and SARS-CoV-2. **B** and **C)** Effects of PSGL-1 on viral release. A PSGL-1 expression vector or control empty vector was cotransfected with the lentivirus packaging construct and a GFP or luciferase reporter plasmid, and viral release was quantified at 72 hours post-transfection by HIV-1 p24 ELISA. **D)** The infectivity of the SARS-CoV pseudotyped virions was quantified by infecting Vero E6 cells and measuring GFP expression at 72 hours post-infection. The percentages of GFP+ cells are shown. **E and F)** The infectivity of the SARS-CoV-2 pseudotyped virions was quantified by infecting Vero E6 (**E**) and Calu-3 cells (**F**). Luciferase activity in infected cells was quantified at 72 hours post infection.

We further tested the effect of PSGL-1 on the infectivity of lentiviral particles pseudotyped with the SARS-Cov-2 S protein. We found that particles pseudotyped with SARS-CoV-2 S protein had much lower infectivity than those pseudotyped with SARS-CoV S protein. To resolve this technical issue, we developed a more sensitive reporter system in which a luciferase reporter (Luc) gene was expressed from the HIV-1 LTR in the presence of co-expressed HIV-1 Tat protein (11) (**Fig. 1A**). A major advantage of this system is that high-level Luc expression can be achieved upon transactivation by co-expressed Tat protein following viral entry, which minimizes non-specific Luc background from non-productive viral entry (12). Using this system, we found that the infectivity of the SARS-CoV-2 pseudovirus is also potently inhibited by the expression of PSGL-1 in the virus-producer cells (**Fig. 1E and 1F**). Together, these results demonstrate that PSGL-1 expression in the virus-producer cells severely diminishes the infectivity of virions bearing SARS coronavirus S proteins.

To investigate possible mechanisms, we analyzed the virion incorporation of SARS-CoV S proteins in the presence of PSGL-1. Our previous study showed that PSGL-1 can inhibit the incorporation of the HIV envelope glycoprotein (1). As shown in **Fig. 2B**, the expression of PSGL-1 in the virus-producer cell also decreased the amount of both SARS-CoV and SARS-CoV-2 S proteins on virions. We and other previously reported that PSGL-1-mediated inhibition of virion infectivity is through steric hindrance of particle attachment to target cells, which does not depend on the presence of viral envelope glycoproteins (1, 5). We performed a virion attachment assay and observed that the lentiviral particles pseudotyped with SARS-CoV or SARS-CoV-2 S protein produced from PSGL-1-expressing cells were impaired in their ability to attach to target cells (**Fig. 2D**). These results demonstrate that the presence of PSGL-1 on virus particles can structurally hinder virion interaction with the target cells even in the presence of remaining S proteins, consistent with previous studies of PSGL-1 and HIV-1 infection (1, 5).

**Fig. 2.**
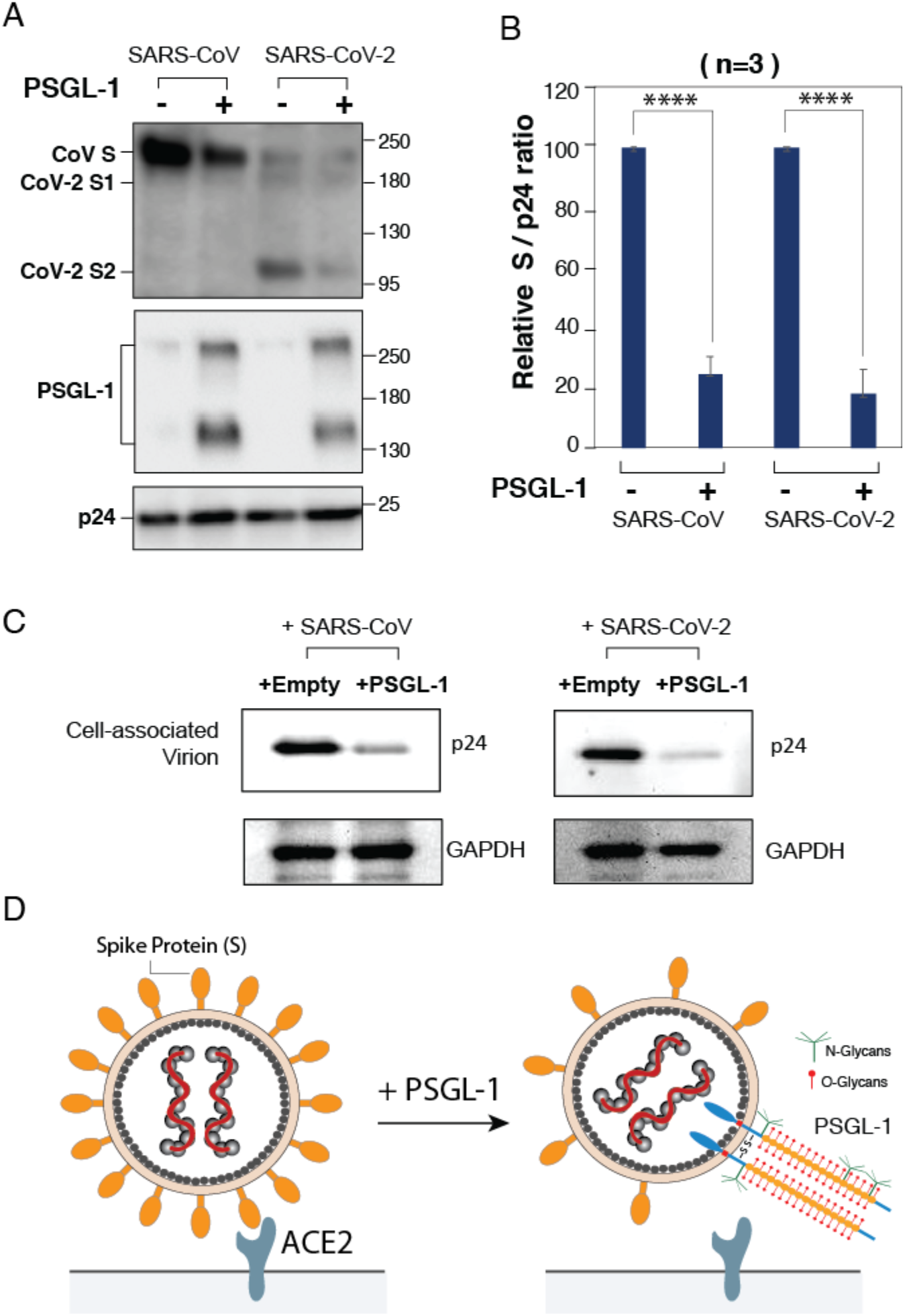
PSGL-1 inhibits virion incorporation of SARS-CoV and SARS-CoV-2 S proteins and blocks virus attachment to target cells. **A**) PSGL-1 inhibits viron incorporation of S proteins. Virions were produced from HEK293T cells cotransfected with HIV-1 pNL4-3/KFS DNA, the vector expressing either the SARS-CoV or the SARS-CoV-2 S protein in the presence of PSGL-1 or an empty control vector. Virion proteins were analyzed by western blotting using antibodies against SARS-CoV S proteins (Genetex), PSGL-1 (KPL-1 clone), or HIV-Ig to detect CA protein p24. **B**) The levels of SARS-CoV or SARS-CoV-2 S proteins in virions were quantified and normalized to viral p24 and set to 100% in the absence of PSGL-1. Data shown are + SD from three independent experiments. P values (two-tailed unpaired t-test): ****p < 0.0001. **C)** PSGL-1 blocks virus attachment to target cells. An equal number of virions produced in the presence of a PSGL-1 vector or empty vector was assayed for attachment to target Vero E6 cells at 4°C for 2 hours. Cells were extensively washed, and cell-associated virions were analyzed by western blot for HIV-1 p24. GAPDH was used as a loading control. **D)** Model for the antiviral activity of PSGL-1 against SARS-CoV and CoV-2 S proteins. Left panel; in the absence of PSGL-1 in the virus-producer cell, virions bearing S protein bind the ACE2 receptor and infect the target cell. Right panel; expression of PSGL-1 in the virus-producer cell results in diminished S protein incorporation, and PSGL-1 incorporation into virions sterically blocks virus binding to target cells.

In this report, we demonstrate that the expression of PSGL-1 in virus-producer cells impairs the infectivity of virions bearing the S protein of either SARS-CoV or SARS-CoV-2, a phenotype shared among several other viruses (e.g., HIV-1, murine leukemia virus, and influenza virus) found to be sensitive to PSGL-1 restriction (1). PSGL-1 has been suggested to be expressed in certain lung cancer cells (13), and in lung phagocytes that control the severity of pneumoccal dissemination from the lung to the bloodsstreem (14, 15). Nevertheless, it remains to be determined whether PSGL-1 is expressed in SARS coronavirus target cells in the lungs, and, if so, whether its expression can impair viral infection. In HIV-1 infection, the viral accessory proteins Vpu and Nef have been shown to antagonize PSGL-1 on CD4 T cells through surface downregulation and intracellular degradation (1, 4). It also remains unknown whether coronaviruses possess a mechanism for antagonizing PSGL-1.

## Materials and Methods

### Virus assembly

The SARS-CoV S and SARS-CoV-2 S protein expression vectors were kindly provided by Gary Whittaker and Nevan Krogan, respectively. A SARS-CoV-2 S protein expression vector was also purchased from Sinobiological. For the production of GFP reporter lentiviral particles pseudotyped with SARS-CoV-S, the SARS-CoV-S expression vector (0.5 μg), pCMVΔR8.2 (7.5 μg), and pLKO.1-puro-TurboGFP (10 μg) were cotransfected with either pCMV3-PSGL-1 (2 μg), or pCMV3-Empty vector (2 μg) as previously described (1). For the production of luciferase reporter lentiviral particles pseudotyped with SARS-CoV-2-S, the SARS-CoV-2-S expression vector (0.5 μg), pCMVΔR8.2 (7.5 μg), and pLTR-Tat-IRES-Luc (10 μg) were cotransfected with either pCMV3-PSGL-1 (2 μg), or pCMV3-Empty vector (2 μg). Both of SARS-CoV-S and SARS-CoV-2-S pseudotyped viral particles were produced in HEK293T cells. Virus supernatants were collected at 48 - 84 hours post transfection, concentrated by ultracentrifugation, and stored at −80°C.

### HIV-1 p24 ELISA

SARS-CoV-S and SARS-CoV-2-S pseudotyped lentiviral particles were quantified using an in-house p24 ELISA kit as previously described (16).

### Virion incorporation of SARS-CoV S proteins

HEK293T cells were co-transfected with HIV-1 Env-defective pNL4-3/KFS (1 μg) and vectors expressing the S protein of either SARS-CoV or SARS-CoV-2 (100 ng) in the presence of PSGL-1 expression vector or an empty control vector (200 ng). Virions were purified and analyzed by SDS-PAGE and western blot using antibodies against SARS-CoV spike proteins (Genetex), PSGL-1 (KPL-1 clone), or HIV-Ig to detect CA protein p24.

### Viral infectivity assay

Virus particles produced in the presence of PSGL-1 or the empty vector were used to infect Vero E6 or Calu-3 cells (ATCC). Cells were pretreated with CoV-2 Pseudovirus Infection Enhancer (CoV-2 PIE), kindly provided by Virongy, for 1 hour at 37°C, and then infected for 5 hours. Infected cells were cultured for 3 days. The percentage of GFP+ cells was quantified by using flow cytometry. For quantifying luciferase activity, cells were washed and lysed in Luciferase Assay Lysis Buffer (Promega). Luminescence was measured by using GloMax^®^ Discover Microplate Reader (Promega).

### Viral attachment assay

Virus particles were incubated with Vero E6 cells (pre-chilled at 4°C for 1 hour) at 4°C for 2 hours. The cells were then washed extensively with cold PBS buffer for 5 times, and then lysed in NuPAGE LDS Sample Buffer (Invitrogen). Cell lysates were analyzed by SDS-PAGE and western blotting with a mouse anti-HIV-1 p24 monoclonal antibody (NIH AIDS Reagent Program, 183-H12-5C) (1:1000 dilution) or anti-GAPDH goat polyclonal antibody (Abcam) (1:1000 dilution) at 4°C overnight, then washed and incubated with anti-mouse IgG, HRP-linked antibody (Cell Signaling) (1:2000 dilution) or anti-goat horseradish peroxidase-conjugated antibody (KPL) (1:2500 dilution) at room temperature for 30-60 minutes. Chemiluminescence signals were developed with Super Signal West Femto Maximum Sensitivity Substrate (Pierce).

## Data availability

All data generated or analyzed during this study are included in this article. Reagents are available from Y. W. upon request.

## Acknowledgements

The authors wish to thank the NIH AIDS Reagent Program for reagents; Drs. Whittaker and Krogan for providing SARS-CoV and SARS-CoV-2 S protein expression vectors, respectively. This work was funded in part by US Public Health Service grants 1R01AI148012 from NIAID to Y.W.. Research in the Freed laboratory is supported by the Intramural Research Program of the Center for Cancer Research, National Cancer Institute, NIH.

## Author contributions

Experiments were designed by Y.W., E.O.F., and K.K.. Manuscript was written by Y.W., and edited by E.O.F. Experiments were performed by S.H., A.A.W., B.H., D.D., and I.V.A..

## Competing interests

A provisional patent applications pertaining to the results presented in this paper has been filed by George Mason University.

## Notes

### Competing Interest Statement

The authors have declared no competing interest.

## References

1. Fu Y, et al. (2020) PSGL-1 restricts HIV-1 infectivity by blocking virus particle attachment to target cells. Proc Natl Acad Sci U S A 117(17):9537–9545.

2. Sako D, et al. (1993) Expression cloning of a functional glycoprotein ligand for P-selectin. Cell 75(6):1179–1186.

3. Somers WS, Tang J, Shaw GD, & Camphausen RT (2000) Insights into the molecular basis of leukocyte tethering and rolling revealed by structures of P- and E-selectin bound to SLe(X) and PSGL-1. Cell 103(3):467–479.

4. Liu Y, et al. (2019) Proteomic profiling of HIV-1 infection of human CD4(+) T cells identifies PSGL-1 as an HIV restriction factor. Nat Microbiol 4(5):813–825.

5. Murakami T, Carmona N, & Ono A (2020) Virion-incorporated PSGL-1 and CD43 inhibit both cell-free infection and transinfection of HIV-1 by preventing virus-cell binding. Proc Natl Acad Sci U S A 117(14):8055–8063.

6. Tang T, Bidon M, Jaimes JA, Whittaker GR, & Daniel S (2020) Coronavirus membrane fusion mechanism offers a potential target for antiviral development. Antiviral Res 178:104792.

7. Raniga K & Liang C (2018) Interferons: Reprogramming the Metabolic Network against Viral Infection. Viruses 10(1).

8. Belouzard S, Chu VC, & Whittaker GR (2009) Activation of the SARS coronavirus spike protein via sequential proteolytic cleavage at two distinct sites. Proc Natl Acad Sci U S A 106(14):5871–5876.

9. Li W, et al. (2003) Angiotensin-converting enzyme 2 is a functional receptor for the SARS coronavirus. Nature 426(6965):450–454.

10. Hoffmann M, et al. (2020) SARS-CoV-2 Cell Entry Depends on ACE2 and TMPRSS2 and Is Blocked by a Clinically Proven Protease Inhibitor. Cell 181(2):271–280 e278.

11. Meltzer B, et al. (2018) Tat controls transcriptional persistence of unintegrated HIV genome in primary human macrophages. Virology 518:241–252.

12. Wang Z, et al. (2010) Development of a nonintegrating Rev-dependent lentiviral vector carrying diphtheria toxin A chain and human TRAF6 to target HIV reservoirs. Gene Ther 17(9):1063–1076.

13. Gong L, Cai Y, Zhou X, & Yang H (2012) Activated platelets interact with lung cancer cells through P-selectin glycoprotein ligand-1. Pathol Oncol Res 18(4):989–996.

14. Tchernychev B, Furie B, & Furie BC (2003) Peritoneal macrophages express both P-selectin and PSGL-1. J Cell Biol 163(5):1145–1155.

15. Ramos-Sevillano E, et al. (2016) PSGL-1 on Leukocytes is a Critical Component of the Host Immune Response against Invasive Pneumococcal Disease. PLoS Pathog 12(3):e1005500.

16. Yoder A, et al. (2008) HIV envelope-CXCR4 signaling activates cofilin to overcome cortical actin restriction in resting CD4 T cells. Cell 134(5):782–792.

